# The paradoxical sustainability of periodic migration and habitat destruction

**DOI:** 10.1101/226589

**Authors:** Zong Xuan Tan, Kang Hao Cheong

## Abstract

Some species and societies engage in sustainable habitat destruction by periodically alternating between a low-growth migratory lifestyle and high-growth but destructive behavior. Examples include nomadic pastoralism and shifting cultivation, practiced by humans for millenia. Although specific models have been developed for species or societies which practice periodic migration and habitat destruction, theoretical insight into such phenomena as a whole is lacking. Here we present a general model of populations which alternate between migratory but negative-growth ‘nomadism’ and destructive ‘colonialism’ which yields high but short-term growth. Despite both strategies individually resulting in extinction, we demonstrate that a population can sustainably colonize an arbitrarily large network of habitats by alternating between the two. This counter-intuitive result can be interpreted in terms of both Parrondo’s paradox and the exploration-exploitation dilemma, suggesting answers to the question of sustainable development.

## 1 Introduction

A number of species, known as ecosystem engineers^1,2^, are capable of significantly transforming the environments they reside in, with humanity itself the archetypal example. Such species alter their habitats in a way that promotes their survival and growth, at least in the short term. In the long term, these alterations can be destructive and unsustainable, as anthropogenic climate change has shown over the past century or so. Yet, there also exist species and societies that survive in stable oscillation with their environments despite these destructive behaviors^3^. One mechanism by which this is possible is a strategy of periodic alternation between destructive but high-growth and non-destructive but low-growth behaviors^4^. For example, many traditional nomadic pastoralists and shifting cultivators are careful to limit the amount of resource depletion that occurs as a result of grazing or farming. By enduring periods of migration to more abundant habitats when their original habitats are sufficiently depleted, they allow the original habitats to recover and remain usable in the future^5,6,7,8^.

While many attempts have been made to model particular instances of the strategy described above, these models have been highly specific to the society or species under study (e.g. army ants, swidden agriculture, nomadic pastoralism)^9,10,11,12,13^. As a result, there remains a lack of theory and insight into this general category of phenomena, and the broad conditions under which survival and growth are successful remain unknown. Optimal foraging models capture some aspects of migratory behavior after a habitat’s resources are depleted^14,15,16,17^. However, because such models assume that organisms forage so as to optimize locally for fitness, they leave out the possibility of switching to low-growth behaviors to ensure long-term survival. Persistence under habitat destruction has been studied using metapopulation approaches, but these approaches assume that destruction is either a random event or externally caused, rather than directly induced by an organism^18,19,20,21^. Models of migratory ecosystem engineers come closest to including all relevant features^22,23^, but, as with other metapopulation approaches, they neither model such engineers as actively destructive of the environment nor as capable of switching behaviors.

Separately, there have been many studies of organisms that alternate behaviors or switch phenotypes to promote resilience and survival, even when the behaviors would individually lead to extinction^24,25^. For example, random phase variation of bacterial phenotypes can ensure subsistence in a temporally varying environments, despite each phenotype being unfit for survival on its own^26^. In the realm of ecology, it has been shown that environmental stochasticity can allow a population can persist by migrating between sink habitats only^27^. Apart from our recent study on the topic^4^, however, there is an absence of research on such counterintuitive ‘reversal behaviors’ in the case of migratory and environmentally destructive species.

As demonstrated in Ref.^4^, the counterintuitive survival of populations which alternate between non-destructive ‘nomadism’ and destructive ‘colonialism’ can be understood as a manifestation of Parrondo’s paradox, which states that there are pairs of losing strategies which can be combined, through alternation, to win^28,29,30,31,32^. There have been many studies exploring the paradox^33,34,35,36,37^. For instance, the evolution of less accurate sensors^38^ and a tumor growth model^39^ have been analysed in terms of the paradox.

In the context of our previous study^4^, ‘nomadism’ was used in a broad sense to refer to any behavioral strategy that has zero or negative growth but also leaves the environment untouched, while ‘colonialism’ referred to strategies which rely on some amount of cooperation and have high rates of short-term growth, but cause environmental destruction in the long run. The former can be seen as analagous to the ‘agitative’ strategy in the original paradox, or Game A, and the latter can be seen as analogous to the history-dependent ‘ratcheting’ strategy, or Game B. More precisely, nomadism is the ‘agitative’ strategy because it allows for environmental resources (analogous to capital in the original paradox) to recover, whereas colonialism is the ‘ratcheting’ strategy because it can exploit an abundance of resources for short-term gains, even though those gains are eventually lost through long-term habitat destruction. Without alternation, both of them result in extinction, but with alternation, colonialism acts as a ‘ratchet’ by periodically exploiting the environmental resources recovered during prior periods of nomadism, thereby ensuring survival.

Although this analysis of what we termed nomadic-colonial alternation captured many aspects of practices like nomadic pastoralism and shifting cultivation, it set aside the highly significant role of inter-habitat migration as part of the nomadic phase of alternation. Such migration can play a crucial role, because it adds an explorative component to nomadism that can counteract the exploitative nature of colonialism — rather than exploit the current environment to the point of no return, populations can migrate to nearby habitats with more resources, enabling not just survival, but population growth. When those habitats are depleted in turn, migration to either the original environment or environments further out can allow the population to continue its growth. Behavioral alternation in the migratory context studied here can thus be seen as not just an expression of Parrodo’s paradox, but also a naturally-occurring solution to the exploration-exploitation dilemma^40,41,42,43^.

To understand the dynamics of this strategy, and the conditions which make it sustainable, we developed and analyzed a general multi-habitat model of nomadic-colonial alternation that incorporates the process of inter-habitat migration. This paper presents the results of that investigation, revealing mathematical and theoretical insights that can apply to multiple real-world systems. Most importantly, it elucidates the conditions under which an environmentally destructive species can sustainably colonize an arbitrarily large network of connected habitats through periodic migration. As an ecological solution to both Parrondo’s paradox and the exploration-exploitation dilemma, the sustainability of this strategy also suggests intriguing possibilities for better addressing what has been called the ‘paradox of sustainable development’^44,45,46^.

## 2 Population model

We model a structured population of individuals spread across n habitat patches, or nodes. Following the nomadic-colonial model developed in our previous work, the population in each habitat *i ≤ n* comprises a sub-population of free-living and migratory nomads, *x_i_*, and a sub-population of environmentally destructive colonists, *y_i_*. Each habitat patch also has an associated carrying capacity *K_i_*, which limits the size of its colonial population.

Within each habitat *i*, colonists are capable of switching to nomadism at a per-capita rate 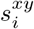, and nomads to colonialism at a per-capita rate 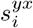. Nomads also migrate from habitat *j* to habitat *i* at a per-capita rate *m_ij_*, whereas colonists stay in their original habitat unless they switch to nomadism. The overall growth rates for the nomadic sub-population *x_i_* and colonial sub-population *y_i_* are thus

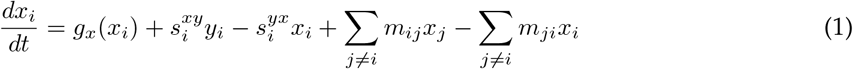

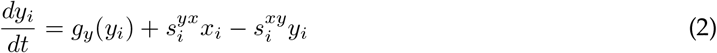

where *g_x_* and *g_y_* are respectively the endogenous nomadic and colonial growth rates (i.e. the growth rates in the absence of both behavioral switching and migration), to be defined below.

### 2.1 Nomadism

Nomads are primarily distinguished from colonists by their ability to migrate to other habitats, as already reflected in Equation 1. They also have a negligible impact on their environment. We restrict our model to the case where pure nomadism leads to extinction in the long run. The endogenous growth rate of the nomadic population *x_i_* in habitat *i* is thus given by

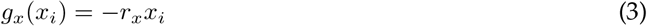

with nomadic decay constant *r_x_* > 0. This restriction is made for two reasons: Firstly, it captures the harsh conditions that nomads often experience while migrating to a new, uncolonized habitat. Secondly, any positive results that are obtained, such as population survival through periodic migration, can then be easily extended to the case where nomadic conditions are more favorable. If survival is ensured under poor conditions, then it can be ensured under better conditions as well.

### 2.2 Colonialism

Colonists are distinguished by the following features: they are subject to both cooperative and competitive effects, and they exploit their environment in order to grow, thereby causing long-term habitat destruction. Cooperation and competition are accounted for in the endogenous growth rate *g_y_* by a modified logistic equation with carrying capacity *K_i_*, Allee capacity *A*, and colonial growth constant *r_y_* > 0:

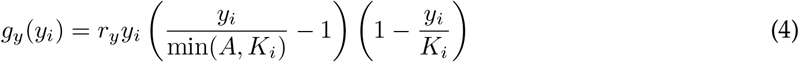

It can be seen that the growth rate is negative when *y_i_* < min(*A*,*K_i_*), a phenomenon known as the strong Allee effect. This captures the necessity of cooperation — colonists need to exceed the critical mass A in order to collectively survive. The growth rate is also negative when *y_i_* exceeds the carrying capacity *K_i_*, due to overcrowding and excessive competition. Positive growth is only achieved when *A* < *y_i_* < *K_i_*, i.e., when the colonial population is neither too large nor too small.

Long-term habitat destruction is accounted for by modelling changes in the carrying capacity *K_i_* as negatively dependent upon the colonial population *y_i_*. Specifically, the rate of change of is given by

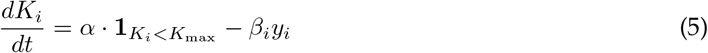

where *α* > 0 is the default growth rate of *K_i_*, *β* > 0 is the per-capita rate of habitat destruction, *K*_max_ is the maximum possible carrying capacity, and **1**_*K_i_* < *K*_max__ is the indicator function which evaluates to one when *K_i_* < *K*_max_, and evaluates to zero otherwise. The indicator function ensures that *K_i_* is limited to a finite maximum of *K*_max_, accounting for the fact that the resources in any single habitat cannot grow infinitely large. This assumption has no effect as long as *K_i_* remains below *K_m_ax*, but serves as a useful simplification of other models which limit the growth of *K_i_* more gradually (e.g., the logistic model presented in Ref. ^4^).

From Equation 5, we can deduce a habitat-stable population level:

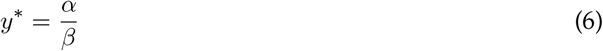

At this population level (*y_i_* = *y^*^*), no habitat destruction occurs 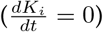. When the colonial population *y_i_* exceeds this level, the carrying capacity decreases, and vice versa. Thus, *y^*^* can also be understood as the long-term carrying capacity of any particular habitat. If this long-term carrying capacity is less than the Allee capacity *A*, the short-term capacity *K_i_* will eventually decrease until it can no longer sustain the critical mass *A* required for colonists to grow. Under these conditions, pure colonialism will be unsustainable in the long run as well.

### 2.3 Behavioral alternation

When should a group of nomads colonize a habitat, and when should the colony then revert to nomadism? A simple and natural rule to follow is to colonize the habitat when resources are abundant, and to switch back to nomadism when resources become depleted, allowing for the exploration of other potential habitats. In accordance with this reasoning, we model the population in each habitat *i* such that it switches to nomadism from colonialism when the carrying capacity is low (*K_i_* < *L*_1_), and switches to colonialism from nomadism when the carrying capacity is high (*K_i_* > *L*_2_). Here, *L*_1_ ≤ *L*_2_ are the switching levels that trigger the alternation of behaviors, assumed to be constant across the entire population. Let *r_s_* > 0 be the switching constant. The colonial-to-nomadic switching rate 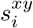 and the nomadic-to-colonial switching rate 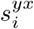 can then be expressed as follows:

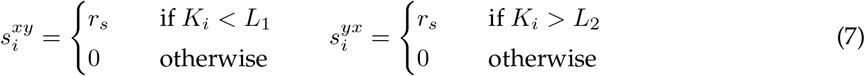

It should be noted that the decision to switch need not always be ‘optimal’ or promote ‘rational’ self-interest (i.e. result in a higher growth rate for each individual). The decision behavior could be genetically programmed or culturally ingrained, such that ‘involuntary’ individual sacrifice promotes the long-term survival of the population.

### 2.4 Reduced parameters

Without loss of generality, we scale all parameters such that *α* = *β* = 1. Equation 5 thus becomes:

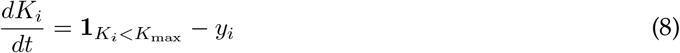

Under this scaling, the habitat-stable population size becomes 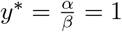, and all other population sizes and capacities are to be interpreted as ratios with respect to *y^*^*. Additionally, since the per-capita rate of habitat destruction *β* =1, *r_x_*, *r_y_* and *r_s_* are to be interpreted as ratios to this rate. As an illustration, if *r_y_* ≫ 1, this means that colonial growth occurs much faster than habitat destruction. Setting *r_x_*, *r_y_* ≫ 1 thus achieves time-scale separation between the population growth dynamics and the habitat change dynamics. Setting *r_s_* ≫ *r_x_*, *r_y_* likewise ensures separation between the dynamics of behavioral switching and population growth.

## 3 Methods

MATLAB R2017a (MathWorks) was used to perform numerical simulations with the included *ode23* ordinary differential equation (ODE) solver. *ode23* implements the Runge-Kutta (2,3) formula pair by Bogacki and Shampine^47^. Accuracy was ensured by repeating each result with consecutively more stringent tolerance levels until the output did not change significantly (i.e. a difference of less than 1%). Both the relative error tolerance and absolute error tolerance were fixed at 10^−6^ after this process.

Exploratory simulations were first conducted for a small number of habitat patches over wide range of parameters and initial conditions. General trends observed from these simulations were then used to guide systematic investigation into the dynamics of migration and colonization. The dynamics of colonization were studied by limiting the initial conditions such that they were progressively less favorable for successful colonization (e.g. by reducing the initial carrying capacities). The observed trends and conditions were then formalized analytically. These conditions were then used to find parameters that ensured survival and expansion for simulations conducted with a large number of habitats.

In deriving these conditions, reasonable assumptions were made in order to make the model analytically tractable. In particular, it was assumed that the rate of behavioral switching was much faster than all other processes (*r_s_* ≫ *r_x_*, *r_y_*, *m_ij_*, 1), and that colonial growth rates were much faster than the rate of habitat destruction (*r_y_* ≫ 1). Initial conditions which result in unstable equilibria (e.g. *y_i_* = *K_i_* = 1 < *A*) were avoided as unrealistic.

## 4 Results

Simulations over a range of parameters showed that the strategy of nomadic-colonial alternation we previously demonstrated to ensure survival in a single habitat was also capable of ensuring survival when extended to multiple habitats. The results also showed that a population localized to a single habitat was capable of colonizing adjacent habitats when resources grew scarce, and then periodically recolonize its original habitat whenever the resources there grew abundant again. Through this strategy of periodic recolonization, a single colony was capable of sustainably expanding to populate all connected habitats in a simulated network, thereby demonstrating that under the right strategy, habitat destruction does not prevent sustainable growth.

Sufficient conditions for the emergence of these phenomena were derived analytically, and are presented in the relevant sections below. As with our work on the single-habitat case, we restrict our results to the case of large switching rates (*r_s_* ≫ *r_x_*, *r_y_*, *m_ij_*, 1), which reflect scenarios where behavior switching can occur more or less instantaneously (i.e. within a fraction of an individual’s lifespan, as is the case for changes in human and animal behavior). This restriction also makes for both conceptual clarity and analytical simplicity, allowing for general insights which can be extended to cases where switching rates are relatively slow as well.

### 4.1 Survival through periodic migration

As noted in our description of the model, neither pure nomadism nor pure colonialism alone can ensure survival when the habitat-stable population level *y*^*^ = 1 is smaller than the Allee capacity *A*. However, as shown in Ref.^4^, a population in a single habitat can ensure survival through nomadic-colonial alternation. Survival was achieved because periodically switching to nomadism allowed the carrying capacity of the habitat to recover after periods of colonial exploitation, and switching to colonialism allowed population levels to recover after periods of nomadic attrition. Unsurprisingly, this finding can be extended to an arbitrary number of isolated habitats (i.e., habitats with no migration between them). Here our results show that *with* the addition of inter-habitat migration, periodic alternation between nomadism and colonialism can ensure survival as well.

Figures 1 and 2 show respectively the survival of populations which periodically migrate between 2 and 3 connected habitats, with equal migration constants *m_ij_* = 5 between all of them. We primarily explain the mechanics with respect to Figure 1 because the dynamics of the two-habitat case can easily be generalized to a larger number of habitats. In Figure 1, habitat 1 is initially populated with colonists, and habitat 2 with nomads. As time passes, it can be seen from Figure 1a that each habitat periodically switches between nomadic and colonial phases — periods of time during which nomadism or colonialism, respectively, are dominant. Furthermore, it can be seen from Figure 1b that the two habitats alternate phases — when nomadism is dominant in habitat 1, colonialism is generally dominant in habitat 2, and vice versa. Hence, when nomads (or colonists) are abundant in habitat 1, they are correspondingly scarce in habitat 2.

**Figure 1:**
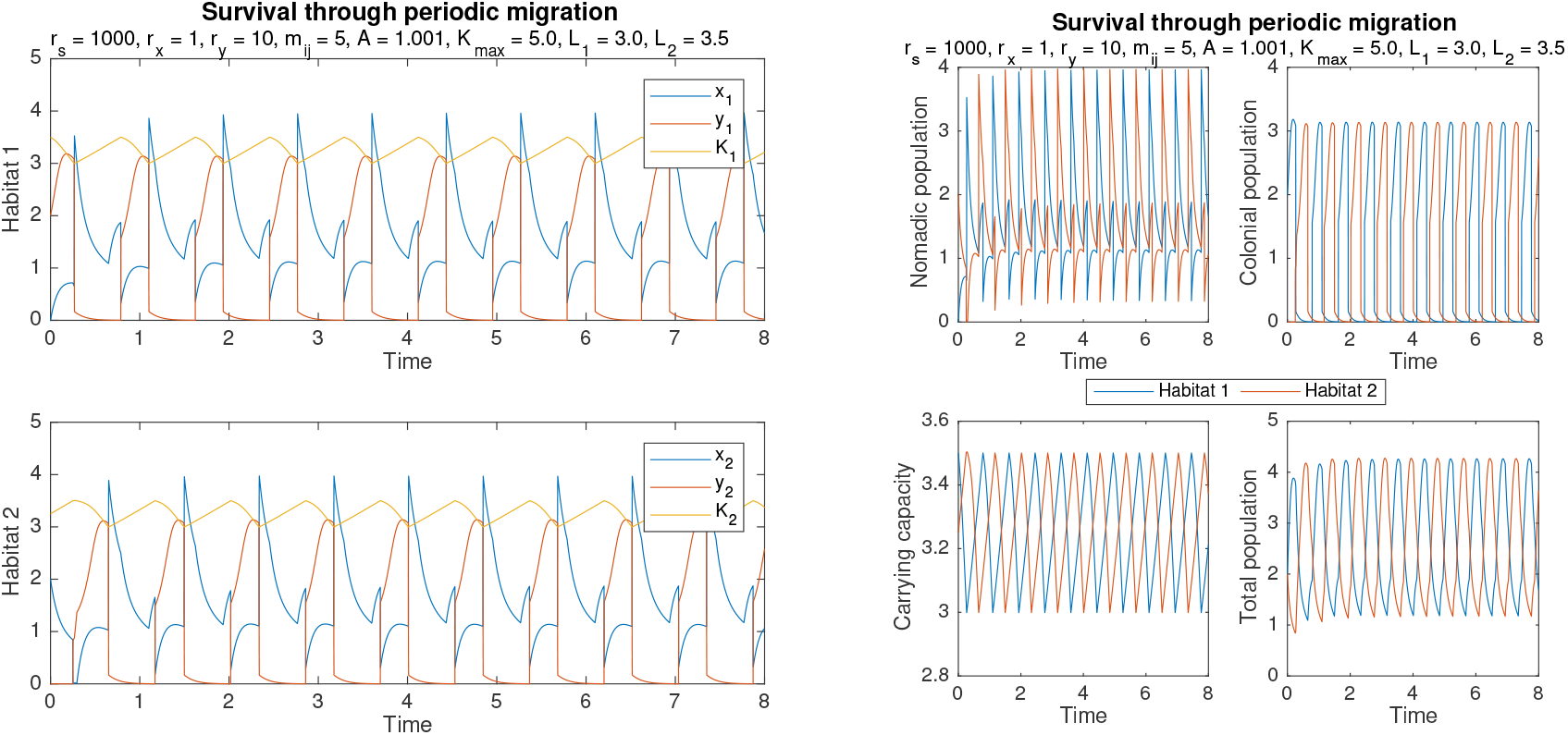
By periodically alternating behaviors and migrating between two habitats, the population ensures its survival. Initial conditions are *x* = [0, 2], *y* = [2, 0], *K* = [3.5, 3.25]. Other parameters are *r_s_* = 1000, *r_x_* = 1, *r_y_* = 10, *∀_i_*, *j*, *m_ij_* = 5, *A* = 1.001, *K_max_* = 5.0, *L*_1_ = 3.0, and *L*_2_ = 3.5

**Figure 2:**
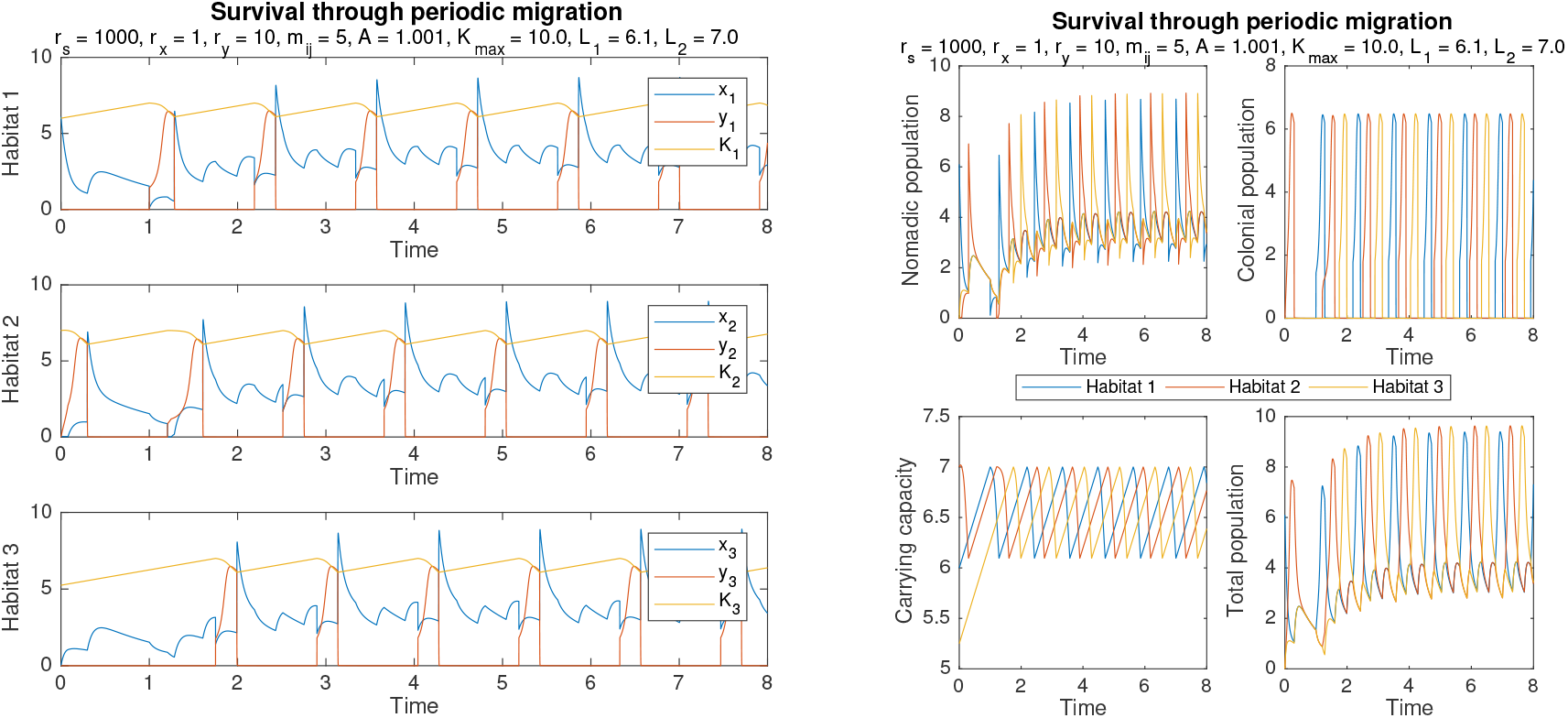
Survival can also be ensured by periodic migration between three habitats, as shown here. Initial conditions are *x* = [6.1,0,0], *y* = [0,0,0], *K* = [6,7,5.25]. Other parameters are *r_s_* = 1000, *r_x_* = 1, *r_y_* = 10, ∀_i_, *j*, *m_ij_* = 5, *A* = 1.001, *K*_max_ = 10.0, *L*_1_ = 6.1, and *L*_2_ = 7.0.

Importantly, when each habitat switches to the nomadic phase (e.g. at *t* ≃ 0.75 in habitat 2), there is a resultant influx of migratory nomads into the adjacent habitat, causing a sudden increase in the nomadic population of that habitat (e.g. *t* ≃ 0.75 in habitat 1). Sometimes a jump in the number of colonists happen instead, because incoming nomads immediately switch behaviors to colonialism (e.g. at *t* ≃ 0.25 in habitat 2). Similar phenomena can be observed in Figure 2, except that in the three-habitat case, several growth spikes occur during each nomadic phase due to migratory influxes from multiple neighbors (e.g. between *t* ≃ 2 and *t* ≃ 3 in the third habitat of Figure 2a).

Survival is achieved for two related reasons. The first is essentially the same as what has been explained for the case of a single habitat. By periodically switching to nomadism, the population in each habitat prevents the resources in that habitat from being depleted, allowing each habitat to be recolonized once resources are abundant again. The second is due to the additional effects of migration. When the population in one habitat switches to nomadism, it then migrates to all adjacent habitats, which then act as a “store” for these nomads. This is particularly effective when those adjacent habitats are near the start of the colonial phase, because the nomads can join the newly-formed colonies and enjoy a period of exponential growth, resulting in a larger population. When this larger population switches back to nomadism, it is better able to recolonize the original habitat once resources there are replenished. Thus, survival is promoted not simply through the strategy of periodic behavioral alternation, but through periodic migration and colonization.

### 4.2 Colonization of unoccupied habitats

Given that the mutual survival of adjacent habitats is enabled by periodic colonization, understanding the dynamics of colonization provides greater insight into the sufficient conditions for survival. In particular, it is useful to examine the colonization of an initially unoccupied habitat, because the conditions which are sufficient for colonizing an unoccupied habitat will also be sufficient for colonizing a habitat which already has a small number of inhabitants.

As an illustration, we analyze the case where a habitat j being colonized is already abundant in resources - i.e., *K_j_* > *L_2_* at the onset of colonization. Prior to onset, habitat *j* is devoid of inhabitants. Figure 3 depicts two such scenarios, where habitat 1 is the source, habitat 2 is the new colony, and *t* = 0 is the onset of colonization. In both scenarios, nomads migrate from habitat 1 to habitat 2, and then switch behaviors from nomadism to colonialism because *K_2_* > *L_2_* = 3.5.

**Figure 3:**
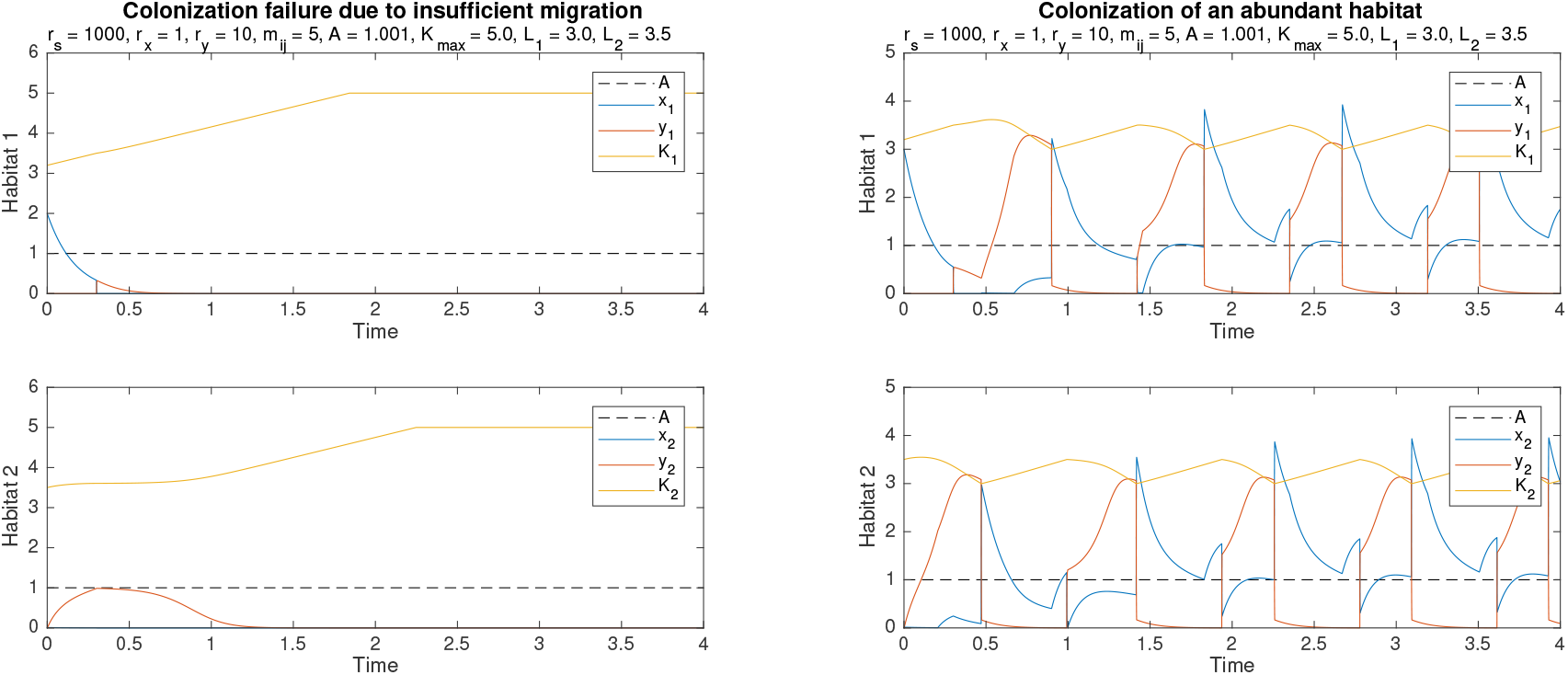
Colonization fails in (a) because of the insufficient number of nomads *x*_1_ in habitat 1 at *t* = 0, but succeeds in (b) because the initial number *x_1_* (0) is higher. Shared initial conditions are *y* = [0, 0], *K* = [3.2, 3.5]. Other parameters are *r_s_* = 1000, *r_x_* = 1, *r_y_* = 10, ∀_i_, *j*, *m_ij_* = 5, *A* = 1.001, *K*_max_ = 5.0, *L*_1_ = 3.0, and *L*_2_ = 3.5.

Colonization ultimately fails in Figure 3a because of the insufficient number of nomads *x_1_* = 2.0 in habitat 1 initially. In Figure 3b however, habitat 1 is initially populated with enough nomads (*x_1_* = 3.0), so the colony in habitat 2 is able to exceed the Allee capacity A due to migration from habitat 1, and from there survive on its own. The population in habitat 2 is then able to re-colonize habitat 1, following which survival through periodic migration ensues.

Multiple factors besides the initial number of nomads influence the success of colonization. Higher rates of nomadic or colonial decay make colonization more difficult, as does a higher Allee capacity A. Rapid migration into a destination habitat makes success more likely, but this has to be balanced against the number of destination habitats that the source population is simultaneously migrating to. If the source tries to colonize too many neighboring habitats at once, not only will it be quickly depleted of nomads without any success. Taking into account all these factors, the following sufficient condition for colonization can be derived (under the assumption of rapid behavioral switching *r_s_* ≫ *r_x_*, *r_y_*, *m_ij_*, 1):

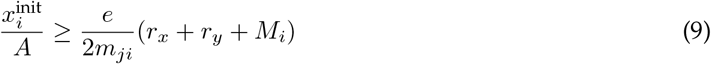

Here *i* is the source habitat, *j* the destination habitat, 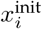 be the initial nomadic population in the source, and 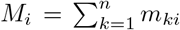 be the total outbound migration constant from *i*. Intuitively, Inequality 9 states that colonization is successful if the ratio of the initial number of nomads 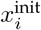 to the Allee capacity *A* exceeds a lower bound defined by the rate parameters.

Analyzing this lower bound gives further insights. Firstly, the bound can be seen to increase with the nomadic decay constant *r_x_*, because more nomadic deaths means less nomads are able to colonize the new habitat. Hence, 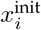 needs to be higher to compensate. The bound also increases with the colonial growth constant *r_y_*. Though counter-intuitive, this is because during colonization, the number of colonists *y_j_* is less than *A*, and so the rate of endogenous nomadic growth *g_y_* is both negative and approximately proportional in magnitude to *r_y_*. Increasing the total outbound migration constant *M_i_* makes the bound higher as well, because more migration means that the initial nomadic population is more quickly depleted. On the other hand, increasing the migration constant *m_ji_* to habitat *j* makes the bound smaller, because more of the migrants go directly to habitat *j* instead of other habitats adjacent to the source.

Other colonization dynamics are possible besides the case just analyzed. Specifically, colonization of both near-abundant and barren habitats can also occur. In general, it is possible to find conditions which allow for successful colonization in all of these cases.

### 4.3 Periodic recolonization of habitats

The above analysis suggests that the initial colonization of habitat *j* is ensured as long as 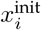 is sufficiently high, but they can also be extended to cover subsequent periods of colonization (e.g., at *t* = 1 in habitat 2 of Figure 3b. As can be seen in habitat 1 of Figure 3b, whenever the source habitat enters the nomadic phase, its nomadic population *x_i_* is at a level of *L*_1_ or above. This occurs because the colonial population *y_i_* quickly grows to reach *K_i_* during the preceding colonial phase. When the switch to nomadism occurs (at which point *y_i_* ≃ *K_i_* ≃ *L*_1_), close to all of the colonists switch behaviors, such that the nomadic population *x_i_* increases almost instantaneously by an amount close to *L*_1_. From this point on, the source habitat *i* has a pool of at least *L*_1_ nomads with which it can re-colonize adjacent habitats. Assuming that the adjacent habitats are abundant at this point in time, we can then replace 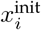 with *L*_1_ in Inequality 9 to obtain conditions for periodic re-colonization:

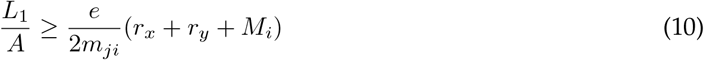

No doubt, there are some circumstances where this specific chain of events does not occur. The colonial growth constant *r_y_* might not be large enough for *y_i_* to reach *K_i_* during the colonial phase, and the adjacent habitats might not always be abundant when the colonial phase begins in the source. However, these circumstances are mitigated by the fact that, once initial colonization of the adjacent habitats has occurred, the total population in those habitats is generally non-zero, making re-colonization easier in the future. Inequality 10 thus serves as a useful guide for finding parameters that result in periodic colonization and survival. Simulation results confirm that satisfying this condition generally produce the desired outcomes.

### 4.4 Sustainable expansion through periodic colonization

With the results derived, it becomes possible to find parameters under which a single population can expand to colonize all connected habitats in an arbitrarily large network. Due to the strategy of periodic behavioral alternation, such outward expansion is sustainable despite the environmentally destructive nature of colonialism. The derived inequalities suggest that expansion is successful across the entire network when the initial population and the switching level *L*_1_ are be sufficiently high. Furthermore, for every pair of connected neighbors *i* and *j*, the migration constant *m_ji_* should be sufficiently large, while the total outbound migration constant *M_i_* should be kept small.

Simulation results demonstrate that sustainable expansion is indeed possible when these considerations are taken into account. Figure 4 shows the population levels over time as a single colony expands to fill a network of *n* = 30 habitats, and Figure 5 shows a visualization of the network as the population spreads from habitat 1. The color bars indicate the corresponding scale, with red representing higher numbers and blue representing lower numbers.

**Figure 4:**
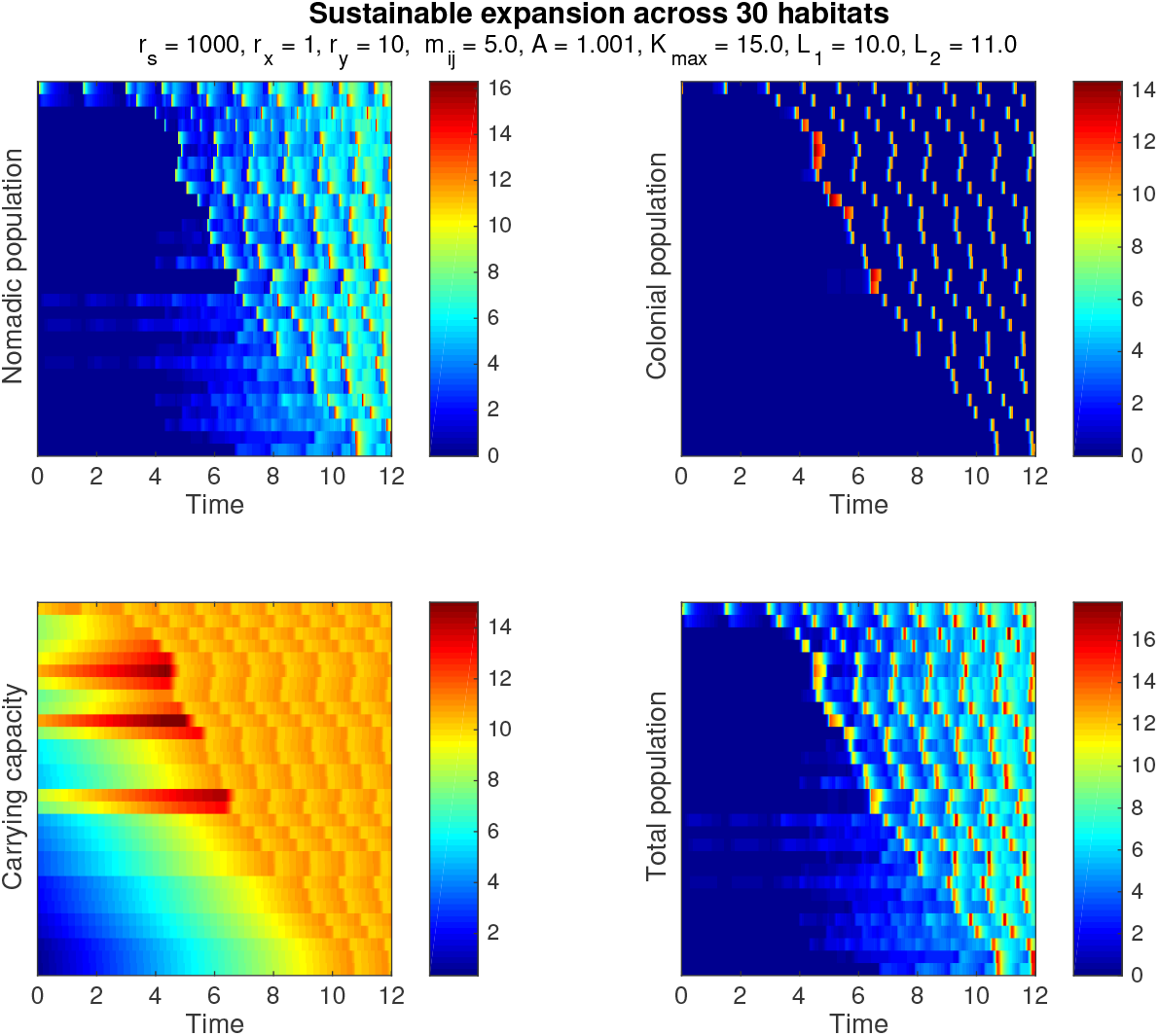
Population and capacity levels over time for a 30-habitat network. Each row of every subplot corresponds to a particular habitat, and the rows are sorted by initial time of colonization (i.e., the time at which *y_i_* exceeds *A* for each habitat *i*). Initial values for the source colony were *x*_1_ = 0, *y*_1_ = 10.5 and *K*_1_ = 11. All other habitats were initially empty (*x_i_* = *y_i_* = 0), with carrying capacities *K_i_* distributed uniformly at random between 0 and 11. Other parameters are *r_s_* = 1000, *r_x_* = 1, *r_y_* = 10, *∀_i_*, *j*, *m_ij_* = 5, *A* = 1.001, *K*_max_ = 15, *L*_1_ = 10, and *L*_2_ = 11.

**Figure 5:**
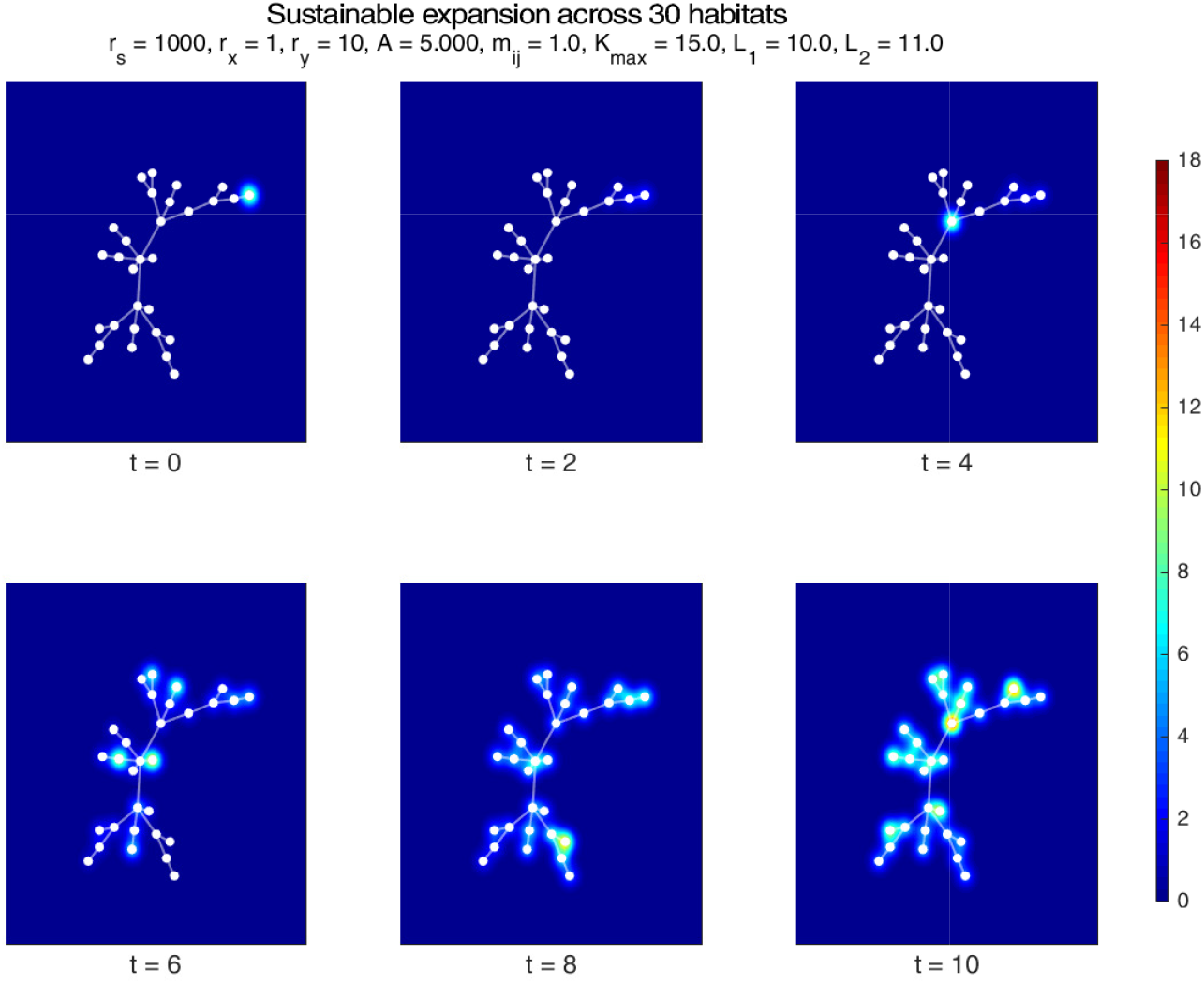
A graphical visualization of the population in Figure 4 spreading across the habitat network, with snapshots taken at various points in time. A 2D Gaussian with a peak value of *x_i_(t)*+*y_i_(t)* is plotted at the corresponding node for each habitat *i*, allowing the total population in each habitat to be visualized.

It can be observed in Figure 5 that the population successfully spreads from the initial colony in the top-right corner of the network to eventually populate all habitats in the network. Figure 4 further shows how this expansion occurs through the same process of periodic migration and colonization described in previous sections. This can be seen most clearly by examining the colonial population levels depicted in the top-right panel of Figure 4. Each row of this panel shows the population levels of a particular habitat, with the rows arranged such that habitats which are colonized first are closer to the top. Initially, only habitat 1 is populated and abundant enough to periodically sustain a population of colonists for short intervals of time. These intervals (i.e. the colonial phases) correspond to the bright orange bars in the first row, whereas all other rows remain deep blue because the habitats they represent are unoccupied. Nearby habitats get colonized by migrants when they grow to have sufficient resources (i.e. *K_i_* > *L_2_*), following which they enter a pattern of periodic migration and recolonization. This can be seen from the bright orange bars that eventually appear in every row.

Similar periodicity can be seen emerging across habitats for both the nomadic population and the carrying capacity. In the case of the nomadic population, the periodicity that can be seen (top-left panel of Figure 4) is less pronounced, since the nomadic phase in each habitat lasts a longer period of time. The periodic alternation of carrying capacity is even less stark (bottom-left panel of Figure 4), because *K_i_* of each habitat *i* just alternates between *L*_1_ = 10 and *L*_2_ = 11 after colonization occurs. Prior to colonization, *K_i_* may increase to values as high as *K*_max_ = 15, before migrants finally arrive and deplete all the excess resources.

The results in Figures 4 and 5 show that expansion is achievable for a network of *n* = 30 habitats under certain initial conditions, but this also extends to networks which are arbitrarily large as long as their maximal degree *d*_max_ is limited. To some degree, expansion was successful in Figures 4 and 5 because habitats nearby the source were sufficiently abundant and could thus be colonized. Nonetheless, parameters can be found such that expansion is also successful under more general circumstances, including cases where barren habitats are present, and where the maximal carrying capacity *K_max_* differs across habitats. In general, sustainable expansion across an arbitrarily large network can be guaranteed over a wide range of circumstances.

## 5 Discussion

By proposing a general model of populations that periodically alternate between explorative nomad-like strategies and exploitative colony-like behaviors, our research provides both mathematical understanding and theoretical unity for a wide range of biological and socio-ecological phenomena. Such phenomena include not just the behavioral alternation of ant colonies^9^ slime moulds^48^, and similar organisms, but also the subsistence strategies of shifting cultivation and nomadic pastoralism that have long been used by humankind.

As explained before, the success of nomadic-colonial alternation as a survival strategy can be understood as a manifestation of Parrondo’s paradox. In the present study however, nomadism is also *explorative* in nature, allowing the population to discover and benefit from new and abundant habitats. This exploration comes at the cost of abandoning a still livable habitat for the harsher conditions of nomadic migration, but it also limits and counteracts the long-term damage of colonialism, which can be understood as *exploitative* in character. Short-term exploitation creates growth that sustains the population as a whole, but over-exploitation of a single habitat makes pure colonialism unviable. Periodic exploration counteracts this not only by limiting the duration of colonialism in any single habitat, but also by finding abundant habitats nearby for the population to exploit instead. By balancing these two processes (thereby satisfying the analytic conditions we have derived), their long-term negative consequences are mitigated, while their short-term positive consequences are preserved. Indeed, appropriately making these trade-offs can be seen as analogous to the exploration-exploitation dilemma^40,41,42,43^. Our results ultimately demonstrate that even when nomadism is so harsh and colonialism so destructive that either alone would lead to extinction in the long run, a population can still spread sustainably across all reachable habitats.

Applying these insights helps explain the success of similar strategies in nature, humanity included. In nomadic pastoralism, groups of pastoralists allow their cattle to graze on fresh pastures until they are somewhat depleted, following which they migrate to find new pastures. In shifting cultivation, agriculturalists cut down areas of forested and fertile land to use as farms until their fertility is reduced, after which they move a new plot of land. Both can be understood as specific forms of the general strategy of alternation modelled here, with periods of grazing or cultivation corresponding to ‘colonialism’, and the migratory periods corresponding to ‘nomadism’. Contrary to common representations and earlier ecological critiques of these practices as ‘unsustainable’, our results show that this is far from necessary. Indeed, there are conditions under which these practices are not only sustainable, but allow for unlimited territorial expansion. This potentially explains why both nomadic pastoralism and shifting cultivation have been so widespread for much of human history. Our model may thus provide a theoretical foundation for the study of these practices in ecological anthropology, similar to how optimal foraging theory has been applied to the study of hunter-gatherer societies^49^. Further analysis of the model could produce new insights about these practices, such as the relationship between the length of fallow periods (i.e. the nomadic phase) and the number of neighboring habitats, or the impact of phase difference between neighboring habitats on total population levels.

By providing an elegant and rigorous framework that can explain a variety of ecological behavior-switching phenomena, our model unifies them into a conceptual whole, allowing general predictions about the chances of survival to be made, and laying the ground for research advances that apply across domains. As a manifestation of Parrondo’s paradox, it also suggests one possible approach to the quite different paradox of ‘sustainable development’. Not all forms of development are sustainable, but perhaps by taking a leaf from the long history of human practices that are environmentally destructive yet paradoxically sustainable, new ways can be found to manage the resources of this planet.

